# The ribosome inhibitor chloramphenicol induces motility deficits in human spermatozoa: a proteomic approach identifies potentially involved proteins

**DOI:** 10.1101/2022.06.28.496361

**Authors:** Marie Bisconti, Baptiste Leroy, Meurig Gallagher, Coralie Senet, Baptiste Martinet, Vanessa Arcolia, Ruddy Wattiez, Jackson Kirkman-Brown, Jean-François Simon, Elise Hennebert

**Author notes:** Corresponding author: Elise Hennebert, Laboratory of Cell Biology, Research Institute for Biosciences, Research Institute for Health sciences and Technology, University of Mons, Place du Parc 23, 7000 Mons, Belgium.

## Abstract

Mature spermatozoa are almost completely devoid of cytoplasm; as such it has long been believed that they do not contain ribosomes and are therefore not capable of synthesising proteins. However, since the 1950s, various studies have shown translational activity within spermatozoa, particularly during their *in vitro* capacitation. Most of them demonstrated that mitochondrial (and not cytoplasmic) ribosomes would be involved in the translation of mitochondrial and nuclear-encoded cytoplasmic mRNAs. However, some evidence suggests that cytoplasmic ribosomes could also be active. Here, we investigate the presence and activity of the two types of ribosomes in mature human spermatozoa. By targeting ribosomal RNAs and proteins, we show that both types of ribosomes are localized in the midpiece as well as in the neck and the base of the head of the spermatozoa. We assessed the impact of cycloheximide (CHX) and chloramphenicol (CP), inhibitors of cytoplasmic and mitochondrial ribosomes, respectively, on different sperm parameters. Neither CHX, nor CP impacted sperm vitality, mitochondrial activity (measured through the ATP content), or capacitation (measured through the content in phosphotyrosines). However, increasing CP concentrations induced a decrease in total and progressive motilities as well as on some kinematic parameters while no effect was observed with CHX. A quantitative proteomic analysis was performed by mass spectrometry in SWATH mode to compare the proteomes of spermatozoa capacitated in the absence or presence of the two ribosome inhibitors. Among the ∼ 700 proteins identified in the different tested conditions, 3, 3 and 25 proteins presented a modified abundance in the presence of 1 and 2 mg/ml of CHX, and 1 mg/ml of CP, respectively. The observed abundance variations of some CP-down regulated proteins were validated using Multiple-Reaction Monitoring (MRM). Taken together, our results show that the sperm motility deficits induced in the presence of CP could be linked to the observed decrease of the abundance of several proteins, at least FUNDC2 and QRICH2.

## Introduction

Translation, the process in which ribosomes synthesize proteins, occurs in all cell types. However, its occurrence in mature spermatozoa has been debated for a long time. During spermiogenesis, which is the final stage of spermatogenesis, the round spermatid develops into a mature motile spermatozoon. During this process the acrosome and the flagellum develop, the DNA in the nucleus undergoes an important compaction, and most of the cytoplasm is ejected (O’Donnell 2014). These two last events have for a long time led scientists to think that mature spermatozoa are transcriptionally and translationally dormant. However, over the past 30 years, many studies have demonstrated the presence of thousands of RNAs, including mRNAs, in spermatozoa (e.g., Chiang et al. 1994, Miller et al. 1999, Ostermeier et al. 2002, Dadoune et al. 2005, Jodar et al. 2013, Sun et al. 2021). It is proposed that mRNAs do not result from direct transcriptional activity but that they would be synthesized during spermatogenesis by spermatogonia, spermatocytes and spermatids and would be stored afterwards in mature spermatozoa (Dadoune et al. 2005, Miller and Ostermeier 2006). Some studies showed that mRNAs stored in spermatozoa are discharged in oocytes during fertilization and could therefore be implicated in early embryogenesis (Ostermeier et al. 2004, Martins and Krawetz 2005, Kumar et al. 2013, Castillo et al. 2018). Additionally, it has been proposed that some mRNAs are translated inside the spermatozoa to support their proper functioning (Gur and Breitbart 2006, Miller and Ostermeier 2006, Zhao et al 2009, Rajamanickam et al. 2017).

The first evidence of translational activities in ejaculated spermatozoa was reported in bulls in the late 1950s, and later in the 1970s in mice and humans, by the incorporation of radiolabelled amino acids into proteins during the incubation of spermatozoa at 37 °C (Bhargava 1957, Prekumar and Bhargava 1972, Mujica 1976, Bragg and Handel 1979). However, these reports stated that protein synthesis was solely mitochondrial (i.e., performed by mitochondrial ribosomes, and involving only mitochondrial genes). Indeed, the incorporation of radiolabelled amino acids was not affected by cycloheximide (CHX), an inhibitor of cytoplasmic ribosome activity, while it was inhibited by chloramphenicol (CP), gentamicin and tetracyclin, which target mitochondrial ribosomes (Prekumar and Bhargava 1972, Mujica 1976, Bragg and Handel 1979). These results were later questioned because the experiments were performed on crude ejaculate, which also contains somatic cells and immature spermatids. It is only in the 2000s that other scientists came to the same conclusion, investigating spermatozoa purified from other cell types and incubated under conditions inducing their capacitation, a process which includes a cascade of physiological changes that spermatozoa must undergo to be able to penetrate and fertilize an oocyte (Gur and Breitbart 2006, Zhao et al. 2009). In addition, some studies showed that nuclear encoded proteins are also synthesized by mitochondrial ribosomes (Gur and Breitbart 2006, Zhao et al. 2009, Rajamanickam et al. 2017). However, as mitochondria use a genetic code which is different from the nuclear one, it is difficult to understand how mitochondrial ribosomes could be able to translate nuclear encoded mRNAs. The mechanism of transport of nuclear-encoded mRNAs from the nucleus to the mitochondria remains also to be elucidated (Amaral et al. 2014a).

Only one published study suggested the potential activity of cytoplasmic ribosomes during capacitation of human spermatozoa, by showing that incorporation of radiolabeled amino acids into proteins is reduced in the presence of CHX (Naz 1998). Moreover, the presence of mono- and polyribosomes has been reported in the sperm cytoplasm, at the level of the neck and the anterior part of the midpiece (Cappallo-Obermann et al. 2011). However, although the sperm proteome contains numerous cytoplasmic ribosomal proteins (Table S1), the presence of complete and functional cytoplasmic ribosomes in mature spermatozoa has been rejected for a long time because intact 28S and 18S ribosomal RNAs (rRNAs) are almost never detected in total RNA extracted from purified spermatozoa (Miller and Ostermeier 2006, Cappallo-Obermann et al. 2011, Johnson et al. 2011).

In view of this literature survey the ability of spermatozoa to produce proteins appears clear, while the type of ribosomes involved remains to be demonstrated. Few studies, of which only two were conducted on human spermatozoa, have identified up-regulated proteins following the capacitation of mammalian spermatozoa (Gur and Breitbart 2006, Zhao et al. 2009, Secciani et al. 2009, Kwon et al. 2014, Rajamanickam et al. 2017, Hou et al. 2019). Interestingly, among the proteins identified in these independent studies, only a small number are recurrent. This discrepancy may be due to the experimental procedures that were used (Western blot quantitation *vs* mass spectrometry). In addition, some focused on CP-inhibited proteins (Gur and Breitbart 2006, Zhao et al. 2009) while others compared non-capacitated and capacitated spermatozoa (Secciani et al. 2009, Kwon et al. 2014, Hou et al. 2019).

In the present study, we investigate the presence and activity of the two types of ribosomes in human spermatozoa. First, we study their localization by targeting their components, *i.e.*, ribosomal RNAs and proteins. Then, we assess the impact of CHX and CP on sperm motility (in terms of both head and tail motion), vitality, mitochondrial activity, and capacitation. Finally, to identify potential translated proteins, we compare the proteome of spermatozoa capacitated in the presence or absence of the two types of ribosome inhibitors using a quantitative proteomic analysis.

## Materials and methods

### Subjects and ethics

Male patients or volunteers aged 18-65 years were recruited for the study. Patients performing a check-up spermogram were recruited at the Ambroise Paré Hospital in Mons (Belgium) whereas the recruitment of volunteers was carried out by poster advertisements on social networks, around the University of Mons (UMons), in the city of Mons or by word of mouth. All experiments conducted in this study were approved by the Ethics Committee of Ambroise Paré Hospital in Mons and by the Ethics Committee of Erasme Hospital in Brussels (protocol P2017/540) and the semen samples were obtained with the informed written consent from all subjects, after a reflection period of at least seven days.

Semen was collected by masturbation after an abstinence period of three to five days, liquified during 15 min and routine seminal analysis was performed according to the World Health Organization (WHO) 2021 guidelines. Only samples whose sperm concentration and motility were within the reference values provided by the WHO guidelines were included in the study.

### Sperm preparation

Purification of spermatozoa from the semen samples was carried out by centrifugation at 300 x g for 20 min at 37 °C on a discontinuous PureSperm 40/80 density gradient (Nidacon) to remove seminal plasma, somatic cells, and immature and dead spermatozoa, as described in Nicholson et al. (2000) and the World Health Organization (WHO) guidelines. Purified spermatozoa recovered from the bottom of the 80% PureSperm fraction were then washed at 600 x g for 10 min at 37 °C with Dulbecco’s phosphate-buffered saline (DPBS). To check the purification efficiency, staining was performed before and after purification using the Diff-Quick kit (RAL Diagnostics). All purified sperm samples contained <1% of potential contaminating cells. Purified spermatozoa were counted on a Makler Chamber and maintained at 37 °C until use.

For all the experiments, spermatozoa were suspended in a capacitation solution composed of HAM’s F-10 Nutrient Mix (31550, Gibco) supplemented with 3 mg/ml HSA (GM501, Gynemed) and 100 µg/ml ampicillin before being processed. This medium was used for two reasons: (1) to prevent sperm aggregation, for immunofluorescence and *in situ* hybridization experiments, and (2) to maintain spermatozoa alive for the duration of the experiments and induce their capacitation, for the study of the influence of ribosome activity inhibitors on sperm parameters and proteome.

### Immunofluorescence

Aliquots of spermatozoa (0.5 x 10^6^) diluted in the capacitation solution were mixed in a 1:1 ratio with 4% paraformaldehyde (PFA) in sodium phosphate buffer (PBS solution, pH 7.4) for 15 min at room temperature, for fixation. The samples were then centrifuged at 2000 x g for 5 min. They were washed twice with 0.05 M glycine in PBS and once with PBS. Then, a total of 0.05 x 10^6^ spermatozoa were spread on 12 mm diameter glass coverslips and air-dried. The spermatozoa were then permeabilized in PBS containing 0.3% Triton X-100 for 20 min and washed in PBS containing 0.05% Tween (PBS-T). The coverslips were incubated in PBS-T containing 3% BSA (PBS-T-BSA) for 30 min and then incubated overnight at 4 °C with rabbit polyclonal anti-RPS6 antibody (2211, Cell Signaling), rabbit polyclonal anti-MRPS27 antibody (17280-AP, Proteintech), or mouse monoclonal anti-RPL3 antibody (FNab07430, FineTest) diluted 1:50, 1:100, and 1:50, respectively, in PBS-T-BSA. Controls were performed by incubating coverslips in PBS-T-BSA without primary antibodies. Following several washes with PBS-T, the coverslips were incubated at room temperature for 1 h with Alexa fluor 568- conjugated goat anti-rabbit (A11011, ThermoFisher Scientific) or anti-mouse (A11004, ThermoFisher Scientific) antibodies diluted 1:100 in PBS-T-BSA. The coverslips were washed 3 times for 5 min with PBS-T and incubated with 60 μg/ml PSA-FITC (FL 1051, Vector Laboratories) in PBS for 30 min in dark at room temperature for acrosome labelling. Finally, the coverslips were washed 3 times in PBS-T and then mounted on microscope slides with Prolong Gold Antifade Mountant with DAPI (P36941, ThermoFisher Scientific). The slides were observed using a confocal microscope Nikon TI2-E-A1RHD25.

### *In situ* hybridization

To localize 28S, 18S, 16S and 12S ribosomal RNAs (rRNAs) in human spermatozoa, specific RNA probes were synthesized from cDNA obtained from HCT116 cells available in the laboratory. This allowed to bypass RNA extraction from spermatozoa, which can be tricky due to the low quantity of RNA in these cells (Pessot et al. 1989, Krawetz 2005). RNA probe synthesis and *in situ* hybridization (ISH) protocols were adapted from Lengerer et al. 2019.

#### RNA extraction and cDNA synthesis

HCT116 cells were cultured in a 6-well plate at 300,000 cells per well in 3 ml McCoy’s 5A medium (Gibco) supplemented with 10% heat inactivated FBS and with 90 UI/mL Penicillin, 90 μG/mL streptomycin for 2 days at 37 °C and 5% CO_2_. After incubation, the culture medium was removed, and the cells were incubated for 10 min at room temperature in 1 ml of TRI Reagent (AM9738, ThermoFisher Scientific). Total RNA was extracted according to ThermoFisher Scientific’s instructions. A 1 μg aliquot of total RNA was submitted to DNase I (1U) for 30 min at 37 °C and reverse transcribed using the qScript cDNA SuperMix (95048-025, QuantaBio) according to manufacturer’s instructions.

#### Probe synthesis

Template DNA for producing DIG-labelled RNA probes were obtained by PCR by using Q5 High-Fidelity DNA polymerase (M0491S, New England Biolabs) with the primers listed in Table S2. Primers were designed with Primer3 (Untergasser et al. 2012). A T7 promoter binding site was added to the reverse strand PCR primers and a Sp6 promoter binding site was added to the forward primers for negative controls. The PCR products were purified using the Wizard SV Gel and PCR Clean-up System (A9281, Promega) and the purified templates were used to produce single stranded digoxigenin (DIG)-labelled RNA probes with the T7 (P2075) and Sp6 (P1085) transcription polymerases from Promega. Transcription was performed after manufacturer’s instructions except for the use of the DIG-labeling mixture (11277073910) from Roche. The RNA probes were diluted at 5 ng/µl in HybMix, composed of 50% formamide, 5 x SSC (0.75 M NaCl, 0.075 M sodium citrate), 100 μg/ml heparin, 0.1% Tween, 0.1% CHAPS, 200 μg/ml yeast tRNA, 1x Denhardt’s, and stored at -80 °C.

#### In situ hybridization

A raw semen sample was washed in capacitation medium and centrifuged at 600 x g for 10 min. The pellet was fixed in 4% PFA in DEPC-treated PBS for 1 h at room temperature. The spermatozoa were then washed twice for 5 min in DEPC-treated PBS containing 0.1% Tween (PBS-T), with centrifugations at 3000 x g for 1 min. They were then dehydrated by ascending methanol series (in PBS-T) and stored at -20 °C in 100% methanol. Spermatozoa were spread on 12 mm diameter glass coverslips and air-dried for 5 min. The coverslips were then transferred to 12 well plates for the following steps. Spermatozoa were rehydrated by a methanol series in PBS-T followed by three washes with PBS-T. Proteinase-K treatment (20 μg/ml in PBS-T) was done at room temperature for 17 min and stopped with Glycine (4 mg/ml in PBS-T). The coverslips were washed 2 x 5 min in PBS-T and incubated 2 x 5 min in 0.1 M TEA, 1 x 5 minutes in 0.1 M TEA with acetic anhydride (400:1), 1 x 5 min in 0.1 M TEA with acetic anhydride (200:1) and 2 x 5 min in PBS-T. Spermatozoa were refixed in 4% PFA in PBS for 20 min at room temperature followed by 5 x 5 min washes in PBS-T. Then, they were heat-fixed at 80 °C for 20 min and incubated in 50% Hybmix in PBS-T at room temperature for 10 min, followed by 2 x 5 min in 100% Hybmix. Coverslips were stored at -20 °C until used. Spermatozoa were prehybridized in fresh HybMix at 55 °C for 2 h. RNA probes were added at a concentration of 0.2 ng/µl after denaturation (7 min at 95 °C and snap chilled on ice). Hybridization was performed for 2 days. The coverslips were then incubated in decreasing Hybmix series in 2x SSC (0.3 M NaCl, 0.03 M sodium citrate) at 62 °C. They were then incubated 2 x 30 min in 2x SSC/0.1% CHAPS at 62 °C for 30 min, followed by 2 x 10 min in MAB (100 mM maleic acid, 150 mM NaCl) at room temperature. Spermatozoa were blocked in 1% blocking solution (11096176001, Roche) in MAB at 4 °C for 2 h. DIG-AP-antibody (11093274910 Roche) incubation was then performed overnight at 4 °C (1:2000 in blocking solution). Spermatozoa were washed 6 x 5 min in MAB at room temperature and were then incubated 2 x 5 min in NTMT (0.1 M NaCl, 0.05 M MgCl_2_, 0.1 M Tris 0.1% Tween-20, pH 9.5). Color development was performed with a NBT/BCIP system (11681451001, Roche) in the dark at 37 °C for 1 h 30 to 2 h 30 according to the RNA probe. For the probes for which a labelling was not observed after 2 h 30, the incubation time was extended to 6 h. Frequent ethanol washes were done to stop the color development, followed by 3 x 5 min washes in PBS-T. Coverslips were finally mounted on microscope slides with 25% glycerol, 10% Mowiol, 0.1M Tris (pH 8.5). Images were taken with a Leica DFC700 T microscope.

### Influence of inhibitors of ribosome activity on sperm parameters

Spermatozoa (3 x 10^6^ cells/ml) were incubated for 4 h in the capacitation medium supplemented or not with different concentrations of cycloheximide (CHX; C7698, Sigma-Aldrich) or chloramphenicol (CP; C0378, Sigma-Aldrich). The two ribosome inhibitors were directly solubilized in the capacitation solution. At the end of the 4 h incubation, the influence of the ribosome inhibitors on different sperm parameters was investigated as follows.

#### 1. Motility

Motility analysis was performed by loading 2 µl of sperm suspension in 10 µm Leja counting chamber slides (SC 10-01-04-B, Microptic) maintained at 37 °C and 5-10 videos (5 sec, 50 fps) corresponding to different fields of the chambers were recorded using a DFK 33UP1300 USB 3.0 color industrial camera connected to an inverted Nikon Eclipse Ts2R Microscope with a 10x negative phase contrast objective. All spermatozoa in each video were analyzed with both Motility Module of the OpenCasa system (Alquézar-Baeta et al. 2019) and FAST software (Flagellar Analysis and Sperm Tracking, Gallagher et al. 2019). OpenCasa was used to analyze the total and progressive motilities. Results were checked manually to avoid counting the same sperm twice. Progressive spermatozoa were differentiated from non-progressive spermatozoa by eye as those swimming actively, either linearly or in a large circle. Where it was unclear whether sperm should be classified as motile or progressively motile, we applied a threshold VSL of 10 um/s (as calculated by OpenCASA) (e.g., Elia et al. 2010). Motility parameters such as curvilinear velocity (VCL), straight line velocity (VSL), average path velocity (VAP), amplitude of lateral head displacement (ALHavg), the power output of the first 30 μm of flagellum (P30), flagellar beat frequency (fBF), and flagellar arcwavelength (fAWL) of progressive spermatozoa (defined as those with track centroid speed TCS > 5 um/s, see Gallagher et al. 2019 for details) were obtained with FAST. About 100 spermatozoa were analyzed per replicate for each condition (see Table S3 for the full number of spermatozoa analyzed).

#### 2. Vitality

A 10 μl aliquot of each sample was mixed with 30 μl of BrightVit solution (Microptic). After 5 min incubation at 37 °C, 25 μl was spread and dried on microscope slides, and mounted with Eukitt (253681.0008, Eurobio scientific). The BrightVit solution is a hypo-osmotic medium that allows the swelling of living cells. The solution is composed of dyes including eosin that penetrates the membranes of dead cells, staining them pink, while living cells remain colorless. In this study, only the hypo-osmotic swelling test was used to determine sperm vitality and 300 spermatozoa were analyzed for each condition.

#### 3. Intracellular ATP

Aliquots of the samples, containing 0.05 x 10^6^ spermatozoa, were mixed with the capacitation solution to a final volume of 100 µl, and transferred into wells of a 96-well plate. A 100 µl aliquot of CellTiter-Glo® 2.0 Reagent (G924A, Promega) was added into the wells, the plate shaken for 2 min to induce cell lysis and then incubated for 10 min at room temperature. The plate was read with a GloMax® Navigator Microplate Luminometer (Promega). The blank-corrected bioluminescence value per condition (sample value minus the value of blank well corresponding to the medium) was calculated for each tested condition. The concentration of ATP (nmol) was calculated by comparison with a standard curve made from a stock solution of 50 mg/ml ATP (A2383, Sigma-Aldrich) in sterile water.

#### 4. Capacitation

The efficiency of capacitation was assessed by phosphotyrosine analysis in Western blot, as tyrosine phosphorylation is recognized as a hallmark for sperm capacitation (Aitken et al. 1996, Naz and Rajesh 2004). Aliquots of the samples, containing 0.4 x 10^6^ spermatozoa, were centrifuged at 2000 x g for 5 min at 4 °C. The spermatozoa were washed 3 times with cold PBS and, after the last wash, the supernatant was removed and the spermatozoa were flash frozen in liquid nitrogen and stored at -80 °C. Proteins were extracted in SDS sample buffer (50 mM Tris, 10% (vol/vol) Glycerol, 2% SDS, 100 mM DTT), heated at 95 °C for 10 min, centrifuged, and loaded on 10% SDS-PAGE gels. After electrophoresis, the proteins were transferred onto PVDF membranes (GE Healthcare) using 25 mM Tris, 192 mM glycine, 0.05% SDS, 20% methanol as transfer buffer. The membranes were washed with PBS containing 0.05% Tween 20 and then blocked for 1 h at room temperature in the same buffer containing 5% BSA. The membranes were incubated overnight with anti-phosphotyrosine clone 4G10 monoclonal antibodies (05-321X, Merk) diluted 1:20,000 in PBS containing 0.05% Tween 20 and 3% BSA. After 5 washes of 5 min in PBS-0.05% Tween 20, HRP-conjugated Goat anti-mouse immunoglobulins (G-21040, ThermoFisher Scientific) diluted 1:100,000 in PBS containing 0.05% Tween 20 and 3% BSA were applied for 1 h. Finally, the membranes were washed again and immunoreactive bands were visualized using the ECL Western Blotting Substrate (32106, ThermoFisher Scientific).

### Sample preparation for mass spectrometry analyses

Spermatozoa (3 x 10^6^ cells/ml) were incubated in the presence or absence of 1 or 2 mg/ml CHX or 1 mg/ml CP for 4 h in the capacitation medium. An amount of 2.5 x 10^6^ spermatozoa was withdrawn to carry out the mass spectrometry (MS) analyses, and the remaining 0.5 x 10^6^ spermatozoa were used to assess capacitation and vitality as described above, to ensure that MS experiments were conducted on samples with similar percentage of capacitated and live spermatozoa.

Spermatozoa were centrifuged at 2000 x g for 5 min at 4 °C and washed 3 times with PBS. The pellets were flash frozen in liquid nitrogen and stored at -80 °C. The samples were resuspended in 50 µl of cold 50 mM K_2_HPO_4_, 8M urea, 50 mM DTT buffer (pH 8.5) and vortexed 3 times 10 sec. Mechanical lysis was performed using an ultrasound probe (IKA U50 sonicator). Three cycles of sonication of 5 sec at 20% amplitude were performed at 4 °C. The samples were centrifuged briefly and incubated for 1 h at room temperature. The sulfhydryl groups of the proteins were then carbamidomethylated with iodoacetamide used in a 2.25-fold excess to DTT in the dark at room temperature for 20 min. The samples were centrifuged at 13,300 rpm for 15 min at 15 °C and the proteins contained in the supernatants were precipitated in cold 80% acetone overnight at -20 °C. After a centrifugation at 13,000 rpm for 20 min at 4°C and acetone evaporation, the resulting pellets were resuspended in 20 μl of 25 mM NH4HCO3 containing 1 μg of modified porcine trypsin (Promega) and incubated for 20 min at 37 °C with agitation (1,300 rpm). They were then incubated overnight at 37 °C without shaking. Trypsinolysis was stopped by adding formic acid to a final concentration of 0.1%. The samples were centrifuged at 13,300 rpm for 15 min and the supernatants were stored at -20 °C.

### Differential proteomic analysis by SWATH mass spectrometry

A quantitative proteomic approach was used to identify differentially regulated proteins between spermatozoa capacitated in the presence or the absence of ribosome inhibitors. Analyses were performed on a UHPLC-HRMS/MS instrument (AB SCIEX LC420 and TripleTOF^TM^ 6600) using SWATH mode of acquisition. Tryptic peptides were separated on a C18 column (YMC-Triat 0.3 mm × 150 mm column) with a linear acetonitrile gradient (5 to 35% of acetonitrile, 5 µl.min^-1^, 75 min) in water containing 0.1% formic acid. MS survey scans (m/z 400-1250, 100 ms accumulation time) were succeeded by 50 SWATH acquisition overlapping windows covering the precursor m/z range. Collision induced dissociation was carried on using rolling collision energy, and fragment ions were accumulated for 50 ms in high sensitivity mode. SWATH technology identifies acquired fragmentation spectra by comparing them to a referential spectral library built through Data-Dependent Acquisition (DDA). As reference, we used a sperm-specific spectral database obtained through a Data Dependant Acquisition (DDA) proteomic analysis performed on proteins extracted from spermatozoa in different conditions using a TripleTof 6600 mass spectrometer (Sciex).

SWATH wiff files were processed using AB SCIEX PeakView 2.2 software and SWATH^TM^ Acquisition MicroApp. Up to six peptides identified with high confidence (>99%) were selected, with six transitions per peptide. Only unshared peptides were subjected to quantification. The XIC extraction window was set to 10 min, and the XIC width was set to 20 ppm. Peptide intensity was calculated as the sum of the area under the curve of the XIC of 6 fragment ions per peptide. The protein intensity was calculated as the sum of the peptide intensities. The protein intensities were extracted and exported in AB SCIEX MarkerView^TM^ 1.2 software which was used for normalization, by dividing each protein abundance by the cumulated protein area of the corresponding sample. Only proteins quantified with a minimum of 2 peptides at a false discovery rate below 1 % were considered.

### Multiple Reaction Monitoring (MRM) analysis of selected proteins

Some of the differentially regulated proteins identified in the SWATH analysis were selected for MRM-based relative quantification. Excepted for two samples, all samples used were obtained from different donors than from the SWATH analysis. The MRM analyses were performed using a QTRAP 6500+ instrument (SCIEX) fitted with an electrospray ionization source (150 °C, 4500V). Test runs were performed on extracted and digested sperm proteins for transition selection and MRM method optimization using the Skyline software (20.2.0.343 MacCoss Lab). Five to six transitions, y or b ions, were chosen for each peptide, and at least two peptides were analyzed for each protein. The same procedure was applied for mitochondrial aconitate hydratase (Q99798), which was used as an internal control. Indeed, its abundance was shown to be stable during capacitation (Castillo et al. 2019). The validated transitions are listed in Table S4. The peptide digests from spermatozoa capacitated in the presence or the absence of ribosome inhibitors were separated on a C18-reversed phase column (YMC TriArt C18, 0,3 mm, 150 mm) and peptides were eluted using a gradient of 5–35% acetonitrile with 0.1% formic acid over 20 min at a flow rate of 5 µl/min. MRM data were acquired in scheduled mode with two minutes retention time window and a maximum cycle time of 1.5 sec. Skyline software (20.2.0.343 MacCoss Lab) was used for visual inspection of MRM data and area under the curve integration. Peak picking for each peptide was manually refined using the transition intensity ratio and retention time as leading parameters. The intensity of all transitions was summed up for each peptide. Protein abundance was obtained as the average of the Ln-transformed area under the curve of each target peptides normalized to the average of the Ln-transformed area under the curve of the aconitate hydratase peptides.

### Statistical analyses

For the analysis of the effect of ribosome inhibitors on sperm parameters, statistical analyses were performed using the python packages statsmodels (v0.13.2, Seabold and Perktold 2010) and pandas (v1.4.2, Reback et al. 2022). For both groups (spermatozoa capacitated in the presence of ribosome inhibitors with respective controls), each parameter of interest was separately fitted to linear mixed effects models to characterise how they varied in response to the concentration of inhibitor. For parameters expressed as percentages, values were logit transformed before fitting. Random effects accounted for donor-dependent responses to inhibitors. The models were specified (e.g. for the curvilinear velocity parameter, VCL) as:

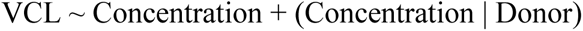

noting that the statsmodels notation suppresses both the fixed and random intercepts. Models were fit using the statsmodels implementation of the Limited-memory Broyden-Fletcher-Goldfarb-Shanno algorithm (Liu and Nocedal 1989). Results were considered statistically significant if p < 0.05 and the model fit converged.

For analysis of the mass spectrometry data, the Shapiro-Wilk test was used to assess assumptions of normality of residuals. Paired t-tests or Wilcoxon tests (in the case of non-normal distribution or if a log transformation did not allow the use of the parametric test) were used to compare the groups using Graph Pad Prism (version 9.0.0, GraphPad software). Results were considered statistically significant if p < 0.05.

## Results

### Localization of ribosomal proteins and RNAs

To investigate the localization of mitochondrial and cytoplasmic ribosomes, we analyzed the presence of ribosomal proteins and RNAs (rRNAs) by immunofluorescence and *in situ* hybridization (ISH), respectively. The labelling obtained for the three proteins investigated was similar, although with slight variations. The mitochondrial ribosome protein MRPS27 was localized at the midpiece, the neck, and the base of the head of the spermatozoa. RPS6, a protein from the small subunit of cytoplasmic ribosomes, was always observed at the level of the neck and the anterior part of the midpiece. Finally, for RPL3, a protein from the large subunit of cytoplasmic ribosomes, we obtained a labelling at the level of the midpiece, the neck and the base of the head (Fig. 1, Figs S1). No labelling was observed in the controls (Fig. S2). In *in situ* hybridization, the staining obtained for the 28S and 18S rRNAs, belonging to cytoplasmic ribosomes, was observed at the level of the base of the head, the neck, and the anterior part of the midpiece of the spermatozoa (Fig. 2, Fig. S3). No labelling was observed for the 12S and 16S mitochondrial rRNAs (Fig. 2) and for the controls (Fig. S4).

**Figure 1.**
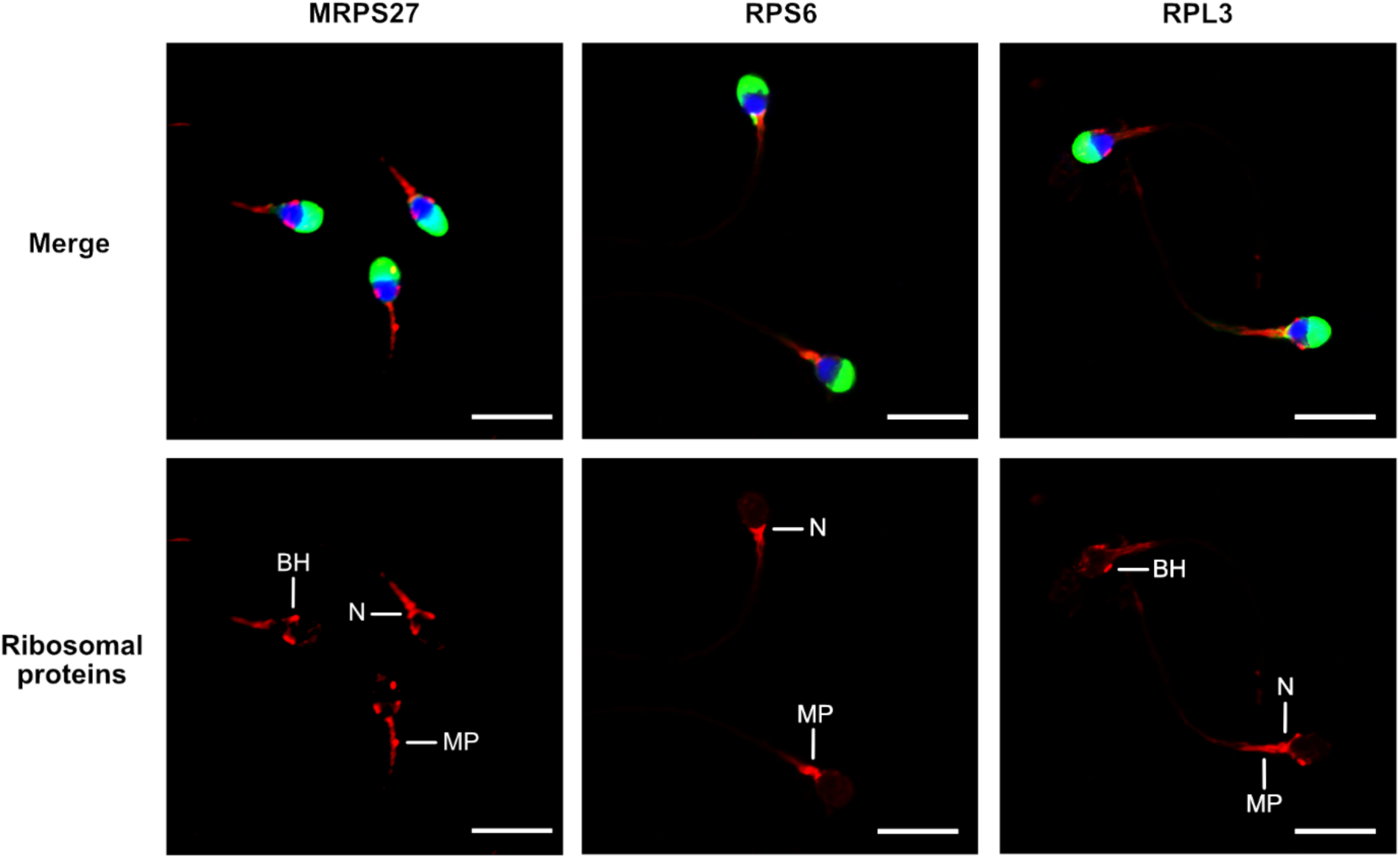
Localization of mitochondrial (MRPS27) and cytoplasmic (RPS6 and RPL3) ribosomal proteins in human spermatozoa. Purified human spermatozoa were fixed with 4% paraformaldehyde, permeabilized with 0.3% Triton-X-100 and stained with anti-MRPS27, -RPS6, or-RPL3 antibodies. Red: Ribosomal proteins, blue: DAPI staining of the nucleus, green: PSA-FITC staining of the acrosome. Scale bar: 10 μm. Representative results of N = 3 experiments. BH: Base of the head, MP: Midpiece, N: Neck.

**Figure 2.**
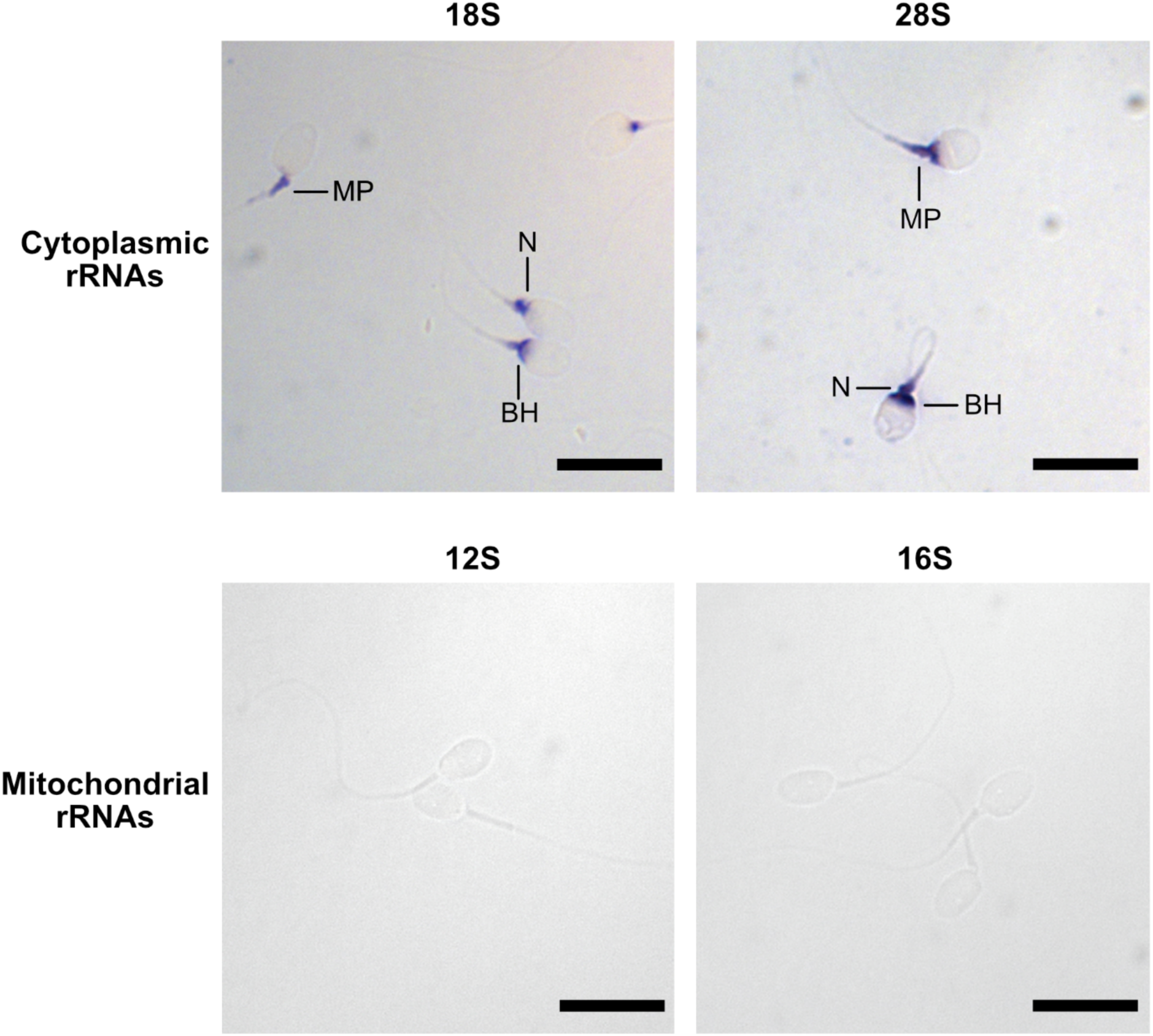
Localization of rRNAs in human spermatozoa by *in situ* hybridization. Spermatozoa were labelled using antisense RNA probes targeting 28S, 18S, 16S, and 12S rRNAs. Scale bar: 10 µm. BH: Base of the Head, MP: Midpiece, N: Neck.

### Influence of inhibitors of ribosome activity on sperm parameters

The relationship between increasing concentrations of cycloheximide (CHX) and chloramphenicol (CP), inhibitors of cytoplasmic and mitochondrial ribosomes, respectively, on different parameters of spermatozoa after 4 h of incubation in a capacitation medium was studied. Regarding sperm motility, while the linear mixed effects models observed no significant relationships with CHX, a reduction in both the total and progressive sperm motility values as CP concentration increases was observed (p = 0.0043 and <0.001 respectively, see Table 1 and Table S5). The percentages of motile and progressive spermatozoa decreased from 75.8% and 69.9% (control) to 66% and 51.9% (CP at 1 mg/ml) respectively, as shown in Figure 3 (A,B). No significant relationships were observed in the presence of increasing concentrations of the two inhibitors for the percentage of live spermatozoa, for ATP content, and for phosphotyrosine content, indicating that the two inhibitors do not impact sperm vitality, the activity of the mitochondria, and sperm capacitation (Table 1, Table S5, Figure 3 C-F).

**Figure 3.**
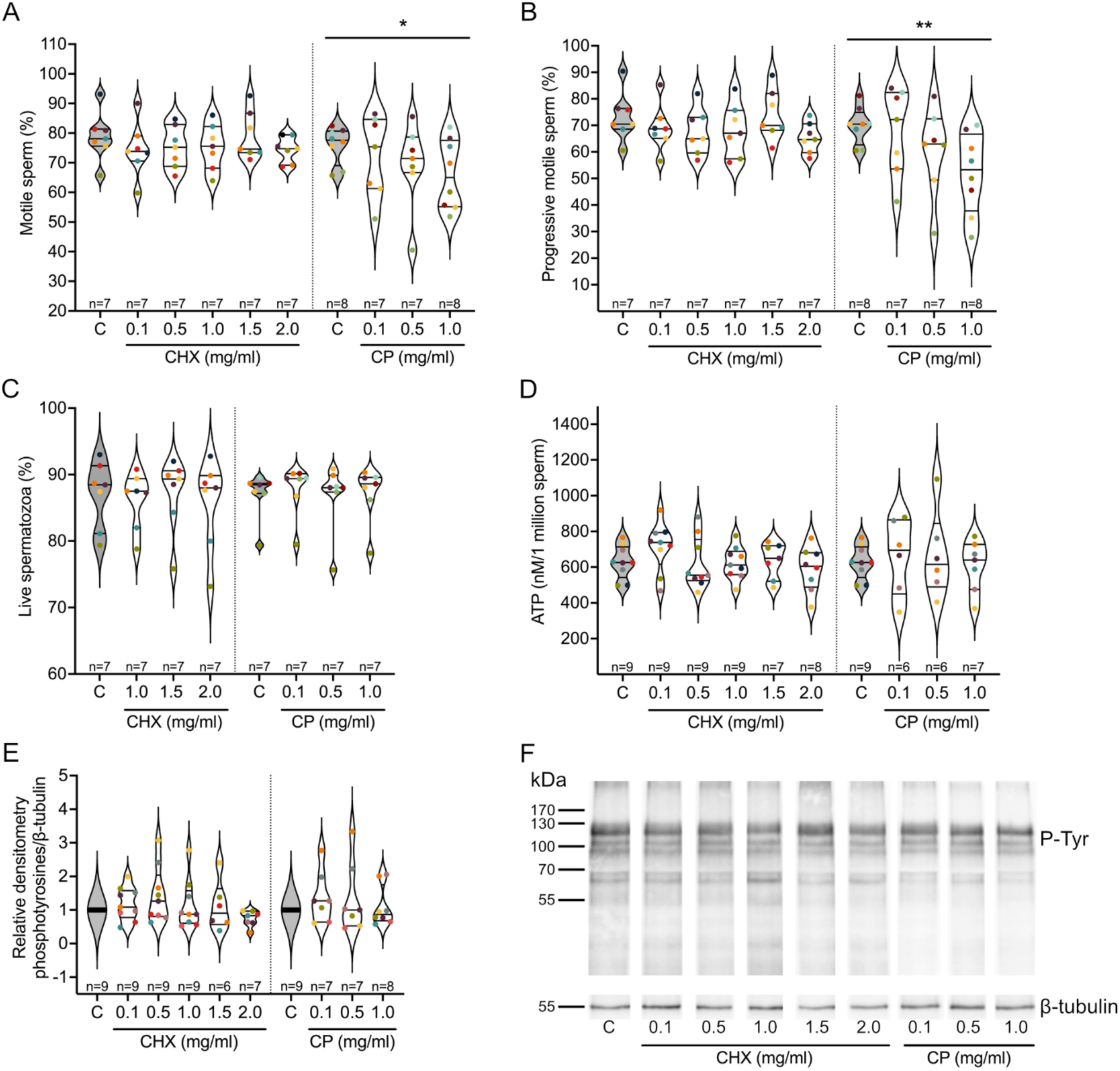
Influence of cytoplasmic (cycloheximide, CHX) and mitochondrial (chloramphenicol, CP) ribosome inhibitors on sperm parameters. Purified human spermatozoa were incubated for 4 h in a capacitation medium in the absence (control, C) or presence of different concentrations of the two types of ribosome inhibitors. (A) Percentage of motile spermatozoa. (B) Percentage of progressive spermatozoa. (C) Percentage of live spermatozoa. (D) ATP content per million of spermatozoa. (E) Quantification of the level of phosphotyrosilated proteins in relative densitometry in respect to β-Tubulin. (F) Representative image of a Western blot analysis of phosphotyrosilated proteins used in (E). Excepted in E, results are presented as violin plots from each condition (i.e., spermatozoa incubated in the presence of different concentrations of ribosome inhibitors). The value for each replicate is highlighted by a dot whose color refers to a donor. * p-value ≤ 0.05, ** p-value ≤ 0.001 (results of the linear mixed-effects models, see Table 1 and Table S5).

**Table 1.**
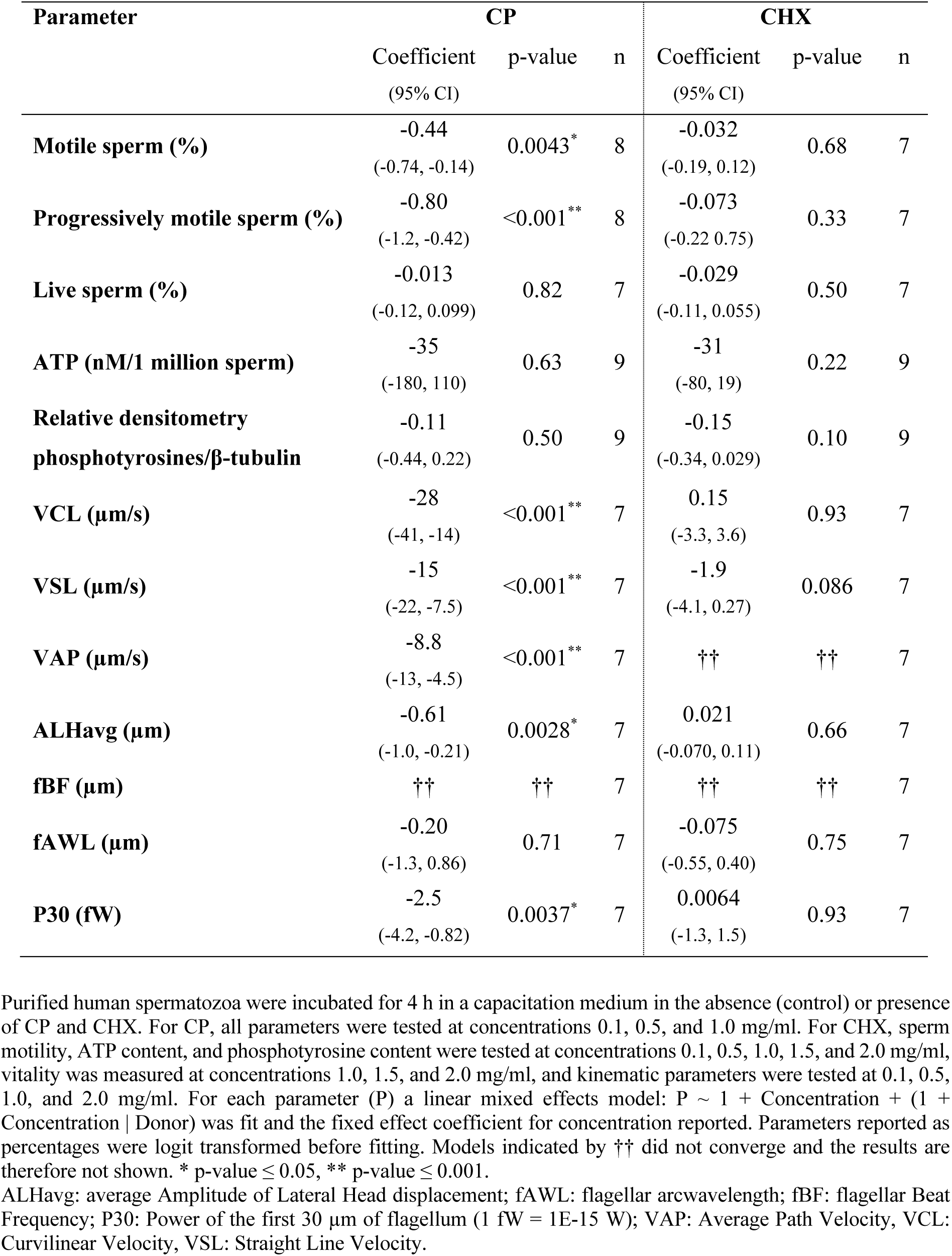
Influence of mitochondrial (chloramphenicol, CP) and cytoplasmic (cycloheximide, CHX) ribosome inhibitors on sperm parameters, shown through fitting linear mixed-effects models. The fixed effect coefficient for concentration of inhibitors are shown with 95% confidence intervals and number of donors n.

| Parameter | CP |  |  | CHX |  |  |
| --- | --- | --- | --- | --- | --- | --- |
|  | Coefficient<br>(95% CI) | p-value | n | Coefficient<br>(95% CI) | p-value | n |
| **Motile sperm (%)** | -0.44<br>(-0.74, -0.14) | 0.0043* | 8 | -0.032<br>(-0.19, 0.12) | 0.68 | 7 |
| **Progressively motile sperm (%)** | -0.80<br>(-1.2, -0.42) | <0.001** | 8 | -0.073<br>(-0.22, 0.075) | 0.33 | 7 |
| **Live sperm (%)** | -0.013<br>(-0.12, 0.099) | 0.82 | 7 | -0.029<br>(-0.11, 0.055) | 0.50 | 7 |
| **ATP (nM/1 million sperm)** | -35<br>(-180, 110) | 0.63 | 9 | -31<br>(-80, 19) | 0.22 | 9 |
| **Relative densitometry**<br>**phosphotyrosines/β-tubulin** | -0.11<br>(-0.44, 0.22) | 0.50 | 9 | -0.15<br>(-0.34, 0.029) | 0.10 | 9 |
| **VCL (μm/s)** | -28<br>(-41, -14) | <0.001** | 7 | 0.15<br>(-3.3, 3.6) | 0.93 | 7 |
| **VSL (μm/s)** | -15<br>(-22, -7.5) | <0.001** | 7 | -1.9<br>(-4.1, 0.27) | 0.086 | 7 |
| **VAP (μm/s)** | -8.8<br>(-13, -4.5) | <0.001** | 7 | †† | †† | 7 |
| **ALHavg (μm)** | -0.61<br>(-1.0, -0.21) | 0.0028* | 7 | 0.021<br>(-0.070, 0.11) | 0.66 | 7 |
| **fBF (μm)** | †† | †† | 7 | †† | †† | 7 |
| **fAWL (μm)** | -0.20<br>(-1.3, 0.86) | 0.71 | 7 | -0.075<br>(-0.55, 0.40) | 0.75 | 7 |
| **P30 (fW)** | -2.5<br>(-4.2, -0.82) | 0.0037* | 7 | 0.0064<br>(-1.3, 1.5) | 0.93 | 7 |

Purified human spermatozoa were incubated for 4 h in a capacitation medium in the absence (control) or presence of CP and CHX. For CP, all parameters were tested at concentrations 0.1, 0.5, and 1.0 mg/ml. For CHX, sperm motility, ATP content, and phosphotyrosine content were tested at concentrations 0.1, 0.5, 1.0, 1.5, and 2.0 mg/ml, vitality was measured at concentrations 1.0, 1.5, and 2.0 mg/ml, and kinematic parameters were tested at 0.1, 0.5, 1.0, and 2.0 mg/ml. For each parameter (P) a linear mixed effects model: $P \sim 1 + \text{Concentration} + (1 + \text{Concentration} | \text{Donor})$ was fit and the fixed effect coefficient for concentration reported. Parameters reported as percentages were logit transformed before fitting. Models indicated by †† did not converge and the results are therefore not shown. \* p-value $\leq 0.05$ , \*\* p-value $\leq 0.001$ .
ALHavg: average Amplitude of Lateral Head displacement; fAWL: flagellar arcwavelength; fBF: flagellar Beat Frequency; P30: Power of the first 30 μm of flagellum (1 fW = $1\text{E-}15$ W); VAP: Average Path Velocity, VCL: Curvilinear Velocity, VSL: Straight Line Velocity.

To further analyze the influence of the inhibitors on sperm motility, we studied different kinematic parameters using FAST. Each of curvilinear velocity (VCL), straight-line velocity (VSL), average path velocity (VAP), amplitude of lateral head displacement (ALHavg) and the power output of the first 30 μm of flagellum (P30) decreased significantly (p < 0.01 for P30 and ALHavg, p < 0.001 for VCL, VSL, VAP) with increasing concentrations of CP, while flagellar arcwavelength (fAWL) had no significant trend and the model fit for flagellar beat frequency (fBF) did not converge. Conversely, CHX did not significantly affect VCL, VSL, ALHavg, fAWL or P30, with the fits for both VAP and fBF failing to converge (Table 1). The full results for the linear mixed effects models are provided in Table S5.

### Influence of inhibitors of ribosome activity on the sperm proteome

A quantitative proteomic analysis was performed using a SWATH approach to compare the proteomes of spermatozoa capacitated in the presence or absence of the two ribosome inhibitors. Based on the results obtained on the sperm parameters, we selected the following conditions: 1 and 2 mg/ml CHX, and 1 mg/ml CP. A total of 688 and 683 common proteins were identified in the different CHX and CP conditions, respectively, with a minimum of 2 peptides. Normalized abundances and fold changes (*i.e.*, ratio of the normalized abundance of a protein in the control condition to its abundance in the tested condition) for all proteins and for each donor are provided in Tables S6 and S7, respectively. Noteworthy, as commercial purified HSA was added in the capacitation medium, we analyzed the proteome of this additive in mass spectrometry and considered that the different proteins identified therein did not belong to spermatozoa. These proteins were not considered for further analyses. We considered that proteins with a mean fold change ≥ 1.25 or ≤ 0.8 and an associated p-value < 0.05 could correspond to potential differentially regulated proteins (Table 2). A very different response was observed depending on the inhibitor tested. Indeed, 25 proteins presented a modified abundance in the presence of 1 mg/ml of CP while, in the CHX conditions, only 3 proteins were significantly differentially regulated in each tested condition. Noteworthy, no protein was common between the two CHX conditions (Table 2).

**Table 2.**
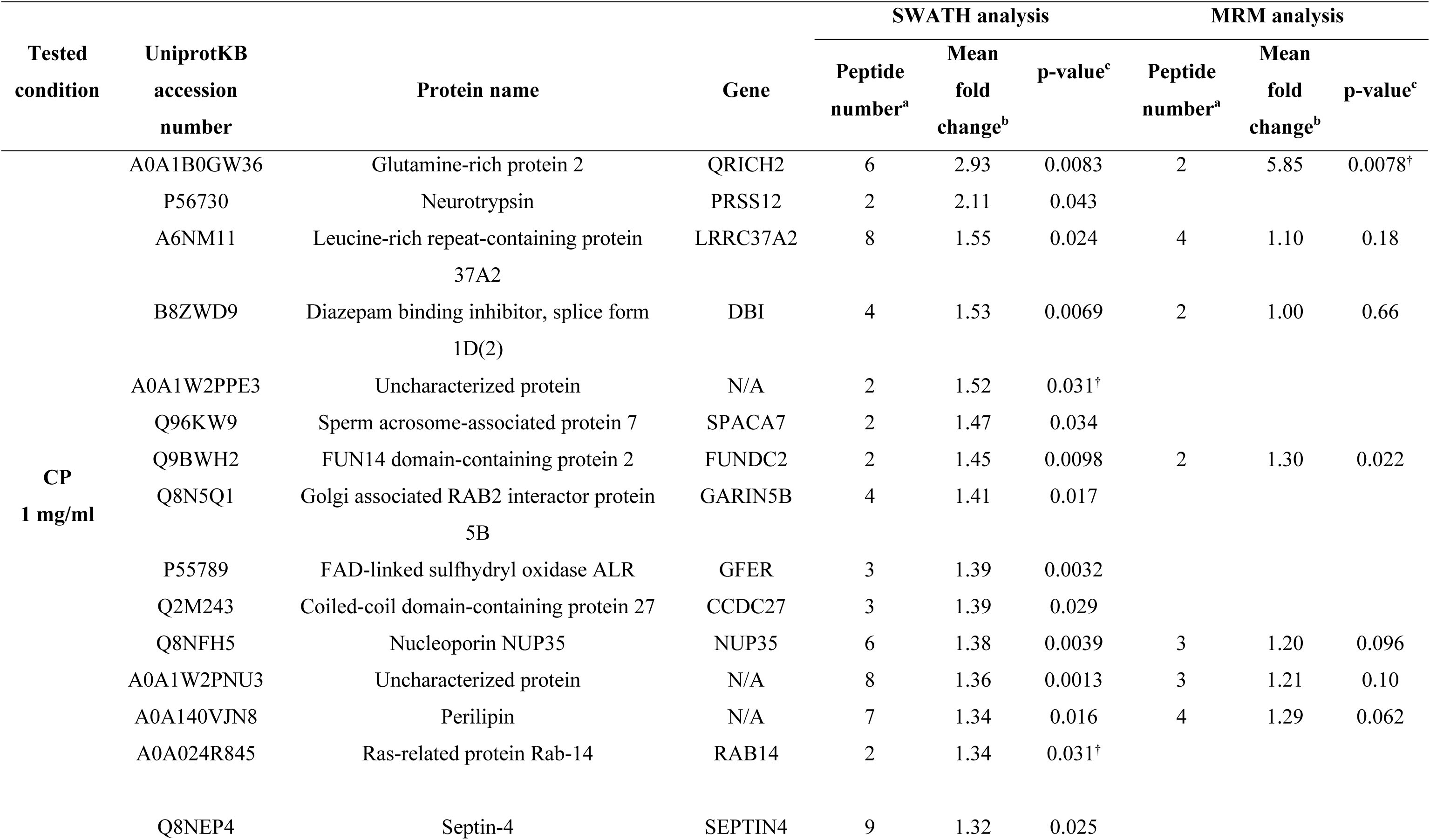
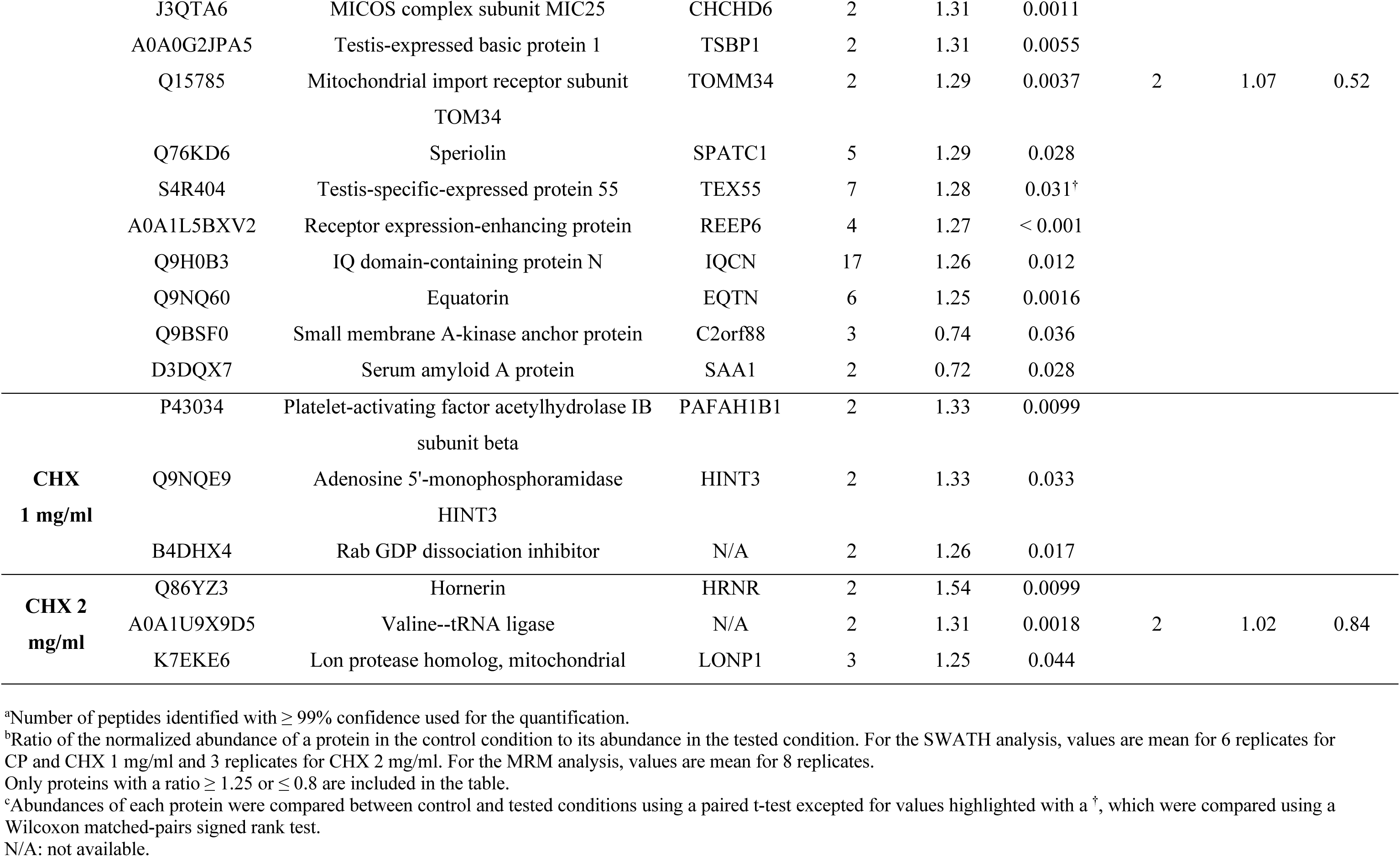
Proteins differentially regulated in spermatozoa capacitated in the presence of chloramphenicol (CP) or cycloheximide (CHX)

To validate these variations, several of the differentially regulated proteins were selected for an MRM-based targeted analysis. We selected these proteins based on the following criteria: i) a stable fold change between different donors (Table S7), and ii) the ability to detect at least 2 peptides with a high-quality MS/MS fragmentation spectrum. Raw data obtained with the MRM analysis are available in Table S8. Among the selected proteins, only QRICH2 and FUNDC2, which were differentially regulated in the SWATH analysis in the CP 1 mg/ml condition, also presented a significant variation in the MRM analysis (Table 2).

## Discussion

Eukaryotic cells have two translation systems: one cytoplasmic, involved in the translation of nuclear encoded proteins by cytoplasmic ribosomes, and one mitochondrial, involved in the translation of proteins encoded in the mitochondrial genome (i.e., 13 polypeptides) by mitochondrial ribosomes (Ott et al. 2016, Wilson and Doudna Cate 2012). In mature spermatozoa, it is well established that the mitochondrial system is active (Bhargava 1957, Prekumar and Bhargava 1972, Mujica 1976, Bragg and Handel 1979, Zhu et al. 2019), while their ability to synthesize nuclear encoded proteins have been debated for a long time. Yet, they contain the machinery involved in cytoplasmic translation, such as ribosomal proteins (Table S1) and translation factors (Wang et al. 2013, Amaral et al. 2014a, Vandenbrouck et al. 2016, Mendonça et al. 2017), as well as mRNAs encoding for them (Sun et al. 2021). Several independent studies showed the up regulation of nuclear encoded proteins following sperm capacitation (Gur and Breitbart 2006, Zhao et al. 2009, Secciani et al. 2009, Kwon et al. 2014, Rajamanickam et al. 2017, Hou et al. 2019) or after sperm exposure to different treatments (e.g., Wang et al. 2018, Kumar et al. 2021). However, the type of ribosomes involved in this process is controverted. Indeed, one study showed the activity of cytoplasmic ribosomes (Naz 1998), while three others claimed the involvement of mitochondrial ribosomes (Gur and Breitbart, 2006; Zhao et al., 2009, Rajamanickam et al. 2017).

In the present study, we contributed to the understanding of the translation process in human spermatozoa by investigating the localization of both types of ribosomes and by studying the influence of ribosome inhibitors on sperm parameters and on the sperm proteome.

### Localization of ribosomes

The presence of complete and functional cytoplasmic ribosomes is questioned in the literature since, during spermiogenesis, the last step of spermatogenesis, most of the sperm cytoplasm is eliminated in the form of “residual bodies” which are phagocytosed by Sertoli cells (Kerr 1992). Moreover, to explain the potential synthesis of nuclear encoded proteins by mitochondrial ribosomes, Gur and Breitbart (2006, 2008) emitted the hypothesis of a possible presence of mitochondrial ribosomes outside the mitochondria. Here, the localization of cytoplasmic and mitochondrial ribosomes was investigated by targeting their proteins and rRNAs.

RPS6 and RPL3, proteins from the small and large subunits of cytoplasmic ribosomes, respectively, were not entirely co-localized in the spermatozoa. RPS6 was found at the level of the neck and the anterior part of the midpiece, a localization consistent with that observed for 28S and 18S rRNAs in ISH. As for RPL3, in addition to the localization observed for RPS6, it was also observed at the level of the whole midpiece and the base of the head. This later localization could correspond to the postacrosomal sheath, a substructure of the perinuclear theca made up of cytosolic proteins (Oko and Sutovsky 2009) and which was recently shown to be enriched in endoplasmic reticulum proteins (Zhang et al. 2022a). The absence of complete co-localization between RPS6 and RPL3 could be explained by the fact that the two ribosome subunits are normally separated in the cytoplasm and only assemble on a mRNA strand, during translation (Hinnebusch and Lorsch 2012). The neck and the anterior part of the midpiece, the common localizations in which the two ribosomal proteins and rRNAs were detected, could therefore presumably contain active cytoplasmic ribosomes. However, the presence of ribosomal proteins and rRNAs in a same site is not a proof of direct interaction and higher resolution techniques, such as immuno-TEM, should be used to confirm this hypothesis. Interestingly, the different labelled regions coincide with the location of the sperm residual cytoplasm, the so-called cytoplasmic droplet, which is predominantly located in the neck and around the midpiece (Bartoov et al. 1980). Our results agree with those obtained by Cappallo-Obermann et al. (2011), who showed that RPS10 and RPL26, proteins from the small and large subunits of cytoplasmic ribosomes, respectively, were localized at the level of the sperm cytoplasm. Interestingly, using TEM, these authors observed ribosomes, and potentially polysomes, in the neck and the anterior part of the midpiece (Cappallo-Obermann et al. 2011). Another study located the ribosomal protein RPL9 in the sperm nucleus by immunofluorescence (de Mateo et al. 2011). Moreover, the same team identified 7 other ribosomal proteins, including RPS6 and RPL3, in the isolated sperm nuclei by mass spectrometry (de Mateo et al. 2011). This localization could be explained by the fact that ribosomal proteins are present in the nucleus during ribosome subunit assembly (Boisvert et al. 2007). While it is obvious that mass spectrometry and immunofluorescence do not provide the same detection sensitivities, it is difficult to explain the completely different localization of RLP9 in the study from de Matteo et al. (2011) and of RPS6 and RPL3 in our study.

It is generally admitted that 28S and 18S rRNAs are degraded in mature human spermatozoa (Cappallo-Obermann et al. 2011, Johnson et al. 2011, Selvaraju et al. 2017, Gòdia et al. 2019, Sellem et al. 2020, Sellem et al. 2021). Therefore, the absence of peaks corresponding to intact rRNAs during the electrophoretic analysis of sperm RNA samples is usually used as a proof that the samples are devoid of contaminating somatic cells (Bianchi et al. 2018, Gòdia et al. 2018). However, one study showed that the density gradient usually used to isolate pure spermatozoa from the semen could be the cause of rRNA degradation (Georgiadis et al. 2015). Here, using ISH, we detected the presence of both cytoplasmic rRNAs in the sperm cytoplasm. However, as we used relatively short probes (∼150 bp), ISH results cannot be a proof of rRNA integrity.

Regarding mitochondrial ribosomes, as expected, MRPS27, from the small subunit, was mainly located in the midpiece, the sperm region containing mitochondria (Challice 1953, Woolley 1970). In addition, the protein was also present in the neck and at the basis of the head (or postacrosomal sheath). This localization could correspond to trace amounts of the protein which was not yet incorporated in mitochondrial ribosomes, or could suggest that mitochondrial ribosomes are also present outside the mitochondria, as observed in *Drosophila* embryos (Amikura et al. 2001). In ISH, we were not able to obtain a labelling for the 12S and 16S rRNAs, while the specificity of the probes was validated on A549 cells (data not shown). This lack of labelling is presumably due to the complex organization of the sperm midpiece, rendering rRNAs difficultly accessible to the probes.

Our results on the localization of cytoplasmic and mitochondrial ribosomes are consistent with those obtained by Gur and Breitbart (2006, 2008), who showed by incorporation of BODIPY-lysine-tRNA that the midpiece was the main site of translation in bovine spermatozoa. Using TEM, they showed that some targeted proteins and corresponding transcripts were localized in the mitochondria (Gur and Breitbart 2006).

### Influence of ribosome inhibitors on sperm parameters

Neither CP nor CHX, inhibitors of the activity of mitochondrial and cytoplasmic ribosomes, appeared to impact the vitality of the spermatozoa, the activity of their mitochondria (measured through the ATP content) and their capacitation (measured through the content in phosphotyrosines) after 4 h incubation in the capacitation medium. These results differ from those obtained by other teams in other organisms. In boar, Zhu et al. (2019) showed a reduced ATP content in spermatozoa incubated for 3 h in CP concentrations ranging from 400 to 800 ng/ml. In their study, they did not incubate the spermatozoa in a capacitation medium but in a low glucose medium, as their aim was to investigate the influence of different energy conditions. As for capacitation, a study performed in mice showed that the percentage of capacitated spermatozoa was decreased after a treatment with 0.1 mg/ml of CP (Zhao et al. 2009). The method used for the detection of capacitated spermatozoa, with Chlortetracycline (CTC) fluorescence assay was different than ours, based on the content of phosphotyrosylated proteins.

Sperm motility was the only investigated parameter impacted during our experiments, with a significant decrease observed using a treatment with CP. Indeed, increasing CP concentrations induced a decrease in total and progressive motilities as well as on the kinematic parameters VCL, VSL, VAP, ALHavg and P30. A similar effect of CP on sperm motility has been observed in boar and bovine spermatozoa, but at lower concentrations (from 400 ng/ml in boar and with 0.1 mg/ml in bull) (Gur and Breitbart 2006, Zhu et al. 2019). To explain the effect of CP on sperm motility, one could hypothesise that CP, through the inhibition of the translation of the 13 proteins encoded in the mitochondrial genome, and involved in the mitochondrial respiratory chain complex, would affect the integrity or functioning of mitochondria, hence inhibiting the production of ATP (Ramalho-Santos et al. 2009, Park and Pang 2021). However, as we did not measure any variation in the sperm ATP content in our experimental conditions, this hypothesis seems unlikely. This leads us to hypothesize that CP affects the translation of proteins important for sperm motility.

### Identification of potentially translated proteins

To identify potentially translated proteins in human spermatozoa, we compared the proteome of spermatozoa capacitated in the presence of different concentrations of both ribosome inhibitors by mass spectrometry in SWATH mode, a label-free technique that allows the quantification of proteins with great robustness (Collins et al. 2017). Based on the results obtained on the sperm parameters, we selected a concentration of 1 mg/ml for CP, as it affected different parameters of sperm motility. As for CHX, as no effect was observed on sperm parameters, we chose to work with 1 and 2 mg/ml, which were the highest tested concentrations and were similar to the concentration tested by Gur and Breitbart (2006) (1 mg/ml) on bovine spermatozoa. A total of 688 and 683 proteins were quantified with a minimum of 2 peptides for the study of the influence of CHX and CP, respectively. This number is comparable to the number of proteins identified with a similar approach in studies comparing different human sperm proteomes (*e.g*., Amaral et al. 2014b, Saraswat et al. 2017, Castillo et al. 2019). As expected, the most abundant identified proteins corresponded to sperm specific proteins (*e.g.*, AKAP4, ODF2) while somatic specific proteins such as E-Cadherin or immunoglobulins were absent. In addition to the use of ampicillin in the samples to remove bacteria, this ensure that the results originate from spermatozoa and not from contaminating cells.

Because it is known that capacitated spermatozoa constitute only a small fraction of the total spermatozoa (Calvo et al. 1993, Cohen-Dayag et al. 1995, Bedu-Addo et al. 2005, Sáez-Espinosa et al. 2020), we considered that proteins with a mean fold change (*i.e.*, ratio of the normalized abundance of a protein in the control condition to its abundance in the tested condition) ≥ 1.25 or ≤ 0.8 and an associated p-value < 0.05 could correspond to potential differentially regulated proteins. With this criteria, 23 proteins were less abundant (fold change ≥ 1.25) in the presence of CP 1 mg/ml compared to the control condition, and 2 proteins were more abundant (fold change ≤ 0.8). Some of the proteins with lower abundance in the CP condition were selected for a validation by MRM-based targeted mass spectrometry on a new cohort of donors. MRM is more sensitive than conventional mass spectrometry and represents a suitable alternative method for the validation of protein abundance variations (Kitteringham et al. 2009, Yocum and Chinnaiyan 2009). Among the eight selected proteins, two, FUN14 domain-containing protein 2 (FUNDC2) and Glutamine-rich protein 2 (QRICH2), presented a confirmed statistically significant lower abundance in the CP condition in this new analysis. Three other proteins, perilipin, nucleoporin NUP35, and an uncharacterized protein appeared to be less abundant in the presence of CP, confirming the trend observed with the SWATH analysis, although the result was not statically significative. And finally, three proteins, leucin-rich repeat-containing protein 37A2 (LRRC37A2), diazepam binding inhibitor (DIB) and mitochondrial import receptor subunit TOM34 (TOMM34) presented a similar abundance between the control and the CP condition, with the MRM analysis. The differences between the SWATH and MRM analyses are presumably due to the very small abundance variations considered in the SWATH analysis. Additionnally, the validation was performed on a different cohort of donors and interindividual variation could also explained the observed differences. Analyzing a larger cohort of samples could help to assess whether the observed trends indeed represent statistically significant differences of abundance.

Interestingly, the two proteins that were validated through the MRM analysis, FUNDC2 and QRICH2, appear to be linked to sperm motility. These results are therefore consistent with the observed sperm motility deficits in the presence of CP. FUNDC2, a component of the outer mitochondrial membrane (Ma et al. 2019), has not been studied much in spermatozoa. However, it has been shown to be down regulated in asthenozoospermic individuals (*i.e.*, presenting low percentage of motile spermatozoa in the semen) (Moscatelli et al. 2019, Yang et al. 2022). QRICH2 is a testis specific protein localized in the sperm tail and involved in its development (Kherraf et al. 2019, Shen et al. 2019, Zhang et al. 2022b). It is also down regulated in asthenozoospermic individuals (Yang et al. 2022).

FUNDC2 and QRICH2 are encoded in the nuclear genome. Their lower abundance in the presence of an inhibitor of mitochondrial ribosomes is consistent with results obtained in other studies showing the impact of CP on nuclear encoded proteins (Gur and Breitbart 2006, Zhao et al. 2009, Rajamanickam et al. 2017). However, these same studies did not identify FUNDC2 and QRICH2 among the deregulated proteins and, in the other way, we did not observe a differential expression of the proteins they highlighted. This discrepancy could presumably be explained by a difference in i) the studied organism (human, bull, mouse or rat), ii) the experimental conditions (e.g., the incubating medium), and iii) the technique used to identify the proteins.

It is noteworthy that the strategy used in the present study does not allow us to affirm that the decrease of abundance of the two proteins is a direct effect of CP on protein synthesis rather than an indirect effect mediated by a regulation of post-translational modifications (PTMs). Indeed, PTMs, by shifting the nominal mass of the modified peptide, induce a decrease in the signal intensity of the non-modified peptide in mass spectrometry. However, at least for QRICH2, this possibility is unlikely, as no common PTMs sites, such as phosphorylation, glycosylation, S-nitrosylation, ubiquitination and SUMOylation, were present in the peptides that were quantified. Moreover, such indirect effect would imply that CP deregulates the enzyme(s) involved in these PTMs. Furthermore, it is not clear from our results whether the synthesis of the identified proteins is induced by capacitation, or on the contrary, is constitutive, to maintain a certain level in the spermatozoa. This question could be answered to by comparing the proteome from non capacitated and capacitated spermatozoa. However, comparing these two conditions is difficult as a lot of biases (*e.g.*, variation in PTMs, presence of HSA or not in the incubating medium, effect on sperm vitality, etc) would influence the results. Actually, Castillo et al. (2019) compared purified (i.e, non capacitated) and capacitated spermatozoa and highlighted the increased abundance of 4 proteins in capacitated spermatozoa: ATP synthase subunit alpha (ATP5F1A), FAM209A, Prohibitin-2 (PHB2) and Tubulin alpha-1A chain (TUBA1A). These proteins were detected in our proteome, but with no abundance variation in the presence of ribosome inhibitors. Finally, the possibility of a non-specific effect of CP mediating the degradation of the differentially regulated proteins cannot be ruled out at this point, we therefore caution our use of the word regulation is to indicate any mechanism influencing abundance.

Among the proteins encoded in the mitochondrial genome (and thus specifically synthesized by mitochondrial ribosomes), only Cytochrome c oxidase subunit 2 (MT-CO2) was identified in our analyses, the others being probably in too low abundance to be detected with the method used. The absence of down regulation of MT-CO2 in the presence of CP while it has been shown that the mitochondrial translation system is active in spermatozoa (Prekumar and Bhargava 1972, Mujica 1976, Bragg and Handel 1979, Zhu et al. 2019) seems to indicate that this protein is not synthesized under our experimental conditions.

Comparatively to CP, CHX did not induce a clear response on the sperm proteome. Indeed, only a few proteins appeared to be differentially regulated under our selected criteria. Moreover, these proteins were not common between the two tested concentrations of CHX, and the only protein which was analyzed in MRM appeared to be stable between the control condition and the 2 mg/ml CHX condition with this method. However, for the SWATH analysis, only 3 replicates were available for the 2 mg/ml CHX condition. That could explain the lack of a robust trend signal. Additional data are needed to validate or not this trend

Using the protocol developed in the present paper, we were only able to analyze a small part of the whole sperm proteome. Indeed, a census of all published sperm proteomes obtained with different techniques described the identification of more than 6000 proteins in human spermatozoa (Amaral et al. 2014a). Therefore, it is likely that the list of differentially regulated proteins in the presence of ribosome inhibitors is incomplete. A better performance for the identification of proteins is often obtained after protein separation by one- or two-dimensional polyacrylamide gel electrophoresis (1D- or 2D-PAGE) (e.g., Amaral et al. 2013, Vandenbrouck et al. 2016). However, differential quantification of proteins with this strategy is less appropriate for the detection of small variations than the gel free strategy we used. The quantity of sperm extract used in the analyses obviously plays a role in the ability to detect proteins in mass spectrometry. Here, this quantity was limited because of the investigation of several conditions per donor. As pointed out above, global proteomic analyses are not always well suited to detect small abundance variations. Another technique, such as MRM should therefore be used to validate the results. Here, we analyzed only few of the proteins differentially regulated in the SWATH analysis in MRM and only two of them could be validated. Therefore, at this point, the differential expression of the other proteins still requires further evaluation.

Overall, our results show that the sperm motility deficits observed in the presence of CP could be linked to the observed decreased abundance of several proteins, at least FUNDC2 and QRICH2. Although other off-target effects of CP on protein abundance cannot be ruled out, a plausible assumption stands in the involvement of mitochondrial ribosomes, which are well documented targets of CP. However, the mechanisms allowing the synthesis of nuclear encoded proteins by mitochondrial ribosomes remain to be elucidated. Even if some studies suggested the presence of mitochondrial ribosomes outside the mitochondria (Gur and Breitbart 2008, Rajamanickam et al. 2017), the difference of genetic codes between mitochondria and nuclei makes this process difficult to understand. Moreover, it is likely that the list of differentially regulated proteins obtained with our strategy is incomplete, and our results do not exclude an activity of cytoplasmic ribosomes. The study of protein translation in spermatozoa undoubtedly offers a new avenue for understanding the molecular mechanisms involved in their function, a prerequisite in the diagnosis and prevention of male infertility.

### Author Contributions

M.B. and E.H. designed research; V.A. and J.-F.S. were involved in donor samples collections; B.L., E.H., C.S., and M.B. performed experimental research; E.H., M.B., B.M., and M.G. performed quantitative and statistical research; B. L., E.H., M.B., B.M., R.W., M.G. and J. K.-B. interpreted the data. E.H. and M.B. drafted the manuscript. All the authors critically reviewed and edited the paper.

### Funding

This research was funded by the Fund for Medical Research in Hainaut (F.R.M.H.), by UMons Research Institute for Biosciences, by UMons Research Institute for Health sciences and Technology (« Kangourou » fund), and by the Fund for Scientific Research of Belgium (F.R.S.- FNRS) under Grant Equipment UN07220F. The Bioprofiling platform used for proteomic analysis was supported by the European Regional Development Fund and the Walloon Region, Belgium. M.B. benefited from a doctoral grant from UMHAP (UMons / Hôpital Ambroise Paré) Medical Research Centre - Scientific Inspiration for Medical Excellence in Mons. B.M. is a Post-doctoral Researcher of the F.R.S.- FNRS. M.G. acknowledges financial support from the University of Birmingham Dynamic Investment Fund.

### Institutional Review Board Statement

The manipulations were conducted according to the guidelines of the Declaration of Helsinki, and by the Ethics Committee of Ambroise Paré Hospital in Mons and by the Ethics Committee of Erasme Hospital in Brussels (protocol P2017/540).

### Informed Consent Statement

Informed consent was obtained from all subjects involved in the manipulations.

## Acknowledgments

We thank P. Quenon from the Laboratory of Cell Biology in UMONS for technical assistance, Dr. C. Decroo from Laboratory of Proteomics and Microbiology in UMONS for his help in mass spectrometry analyses, Pr. D. Nonclercq from the Laboratory of Histology in UMONS for the use of his Leica microscope, all the technicians of the fertility clinic of the Ambroise Paré Hospital, especially C. Barthe, for their help in the recruitment of patients and voluntary donors, and Prof. D. Smith from the University of Birmingham for useful discussions regarding the statistical analysis.

