## Supplementary tables and figures for "The ribosome inhibitor chloramphenicol induces motility deficits in human spermatozoa: a proteomic approach identifies potentially involved proteins"

***Corresponding author:**

### Supplementary Figures

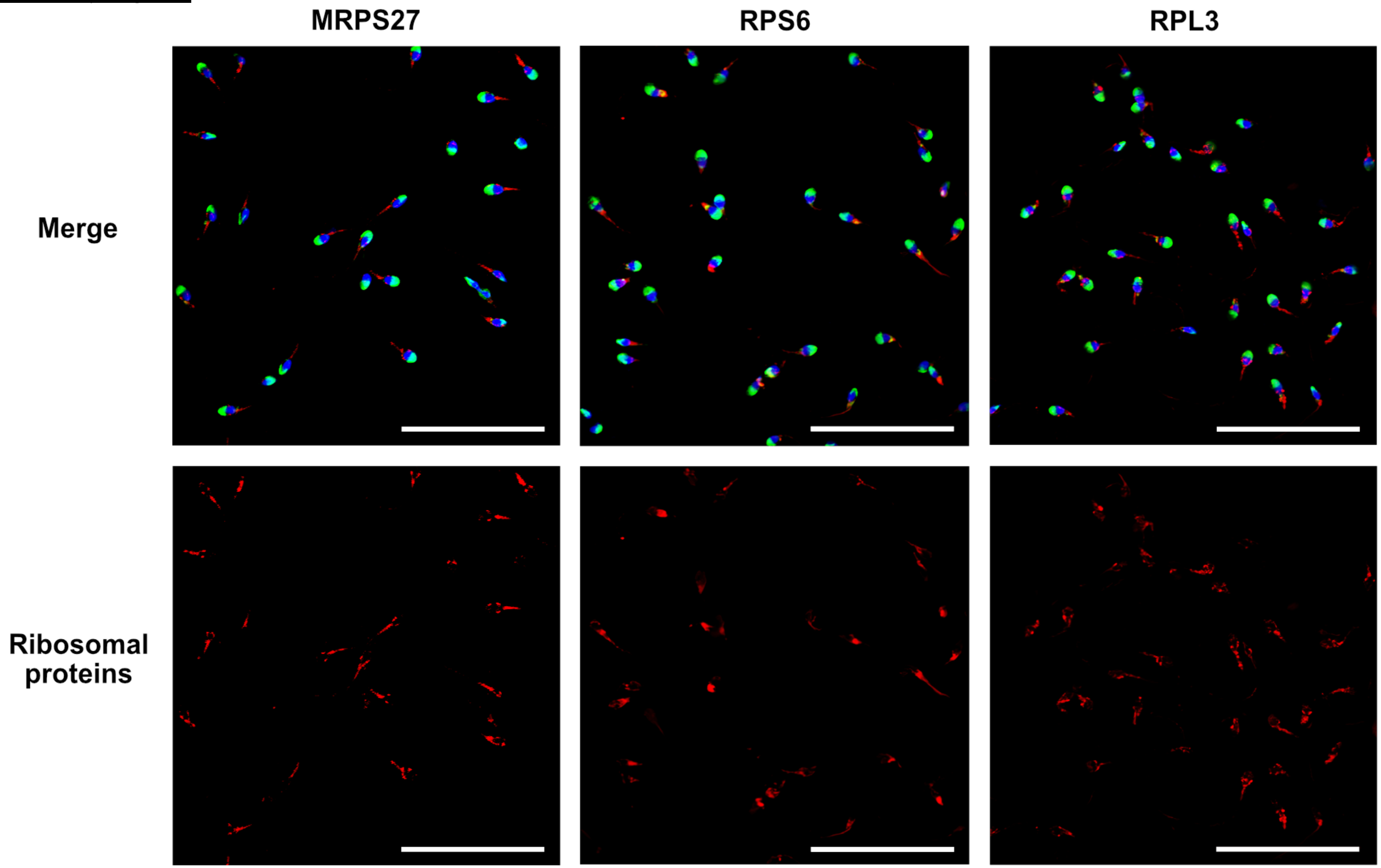


**Figure S1**. General views for the localization of mitochondrial (MRPS27) and cytoplasmic (RPS6 and RPL3) ribosomal proteins in human spermatozoa. Purified human spermatozoa were fixed with 4% paraformaldehyde, permeabilized with 0.3% Triton-X-100 and stained with anti-MRPS27, -RPS6, or-RPL3 (antibodies. Red: Ribosomal proteins, blue: DAPI staining of the nucleus, green: PSA-FITC staining of the acrosome. Scale bar: 50 µm. Representative results of N = 3 experiments.

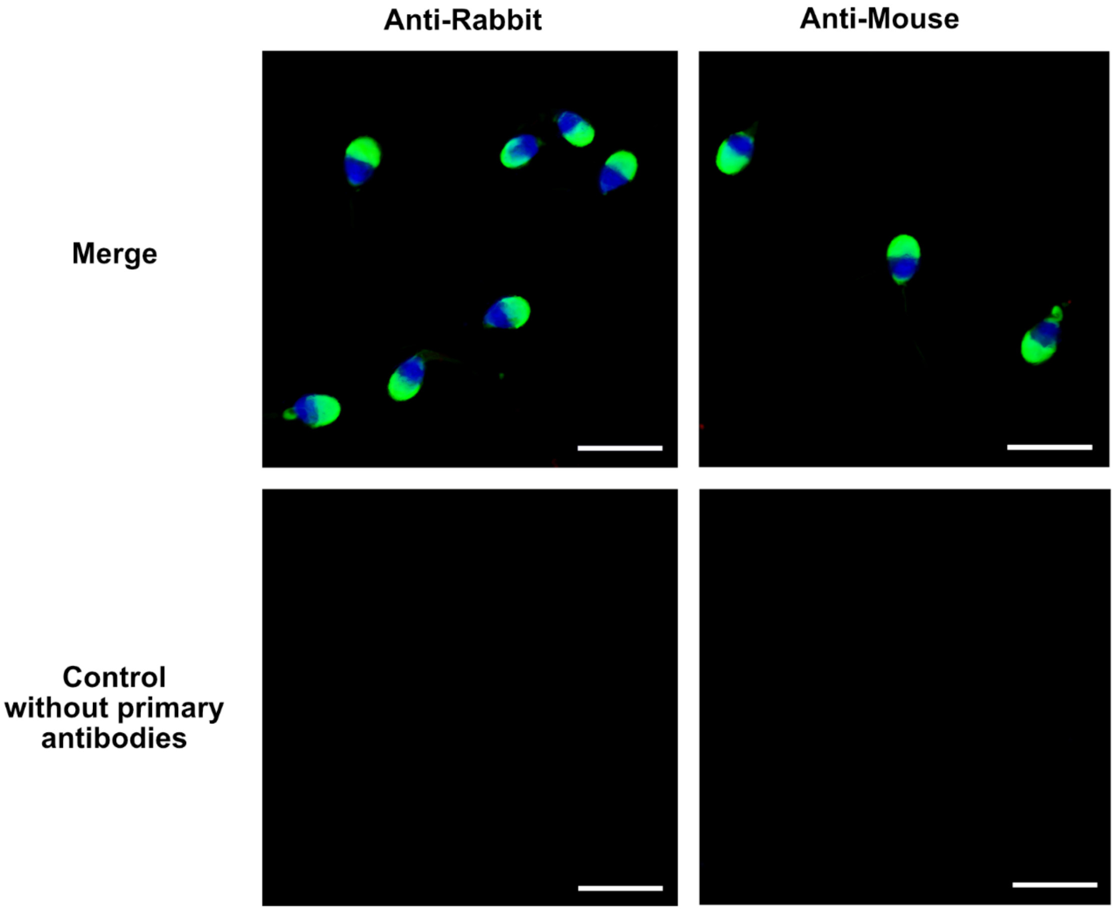


**Figure S2.** Negative controls for the localization of mitochondrial and cytoplasmic ribosomal proteins in human spermatozoa. Purified human spermatozoa were fixed with 4% paraformaldehyde, permeabilized with 0.3% Triton-X-100 and directly incubated with Alexa fluor 568-conjugated goat anti-rabbit or anti-mouse antibodies. Blue: DAPI staining of the nucleus, green: PSA-FITC staining of the acrosome. Scale bar: 10 µm. Representative results of N = 3 experiments.

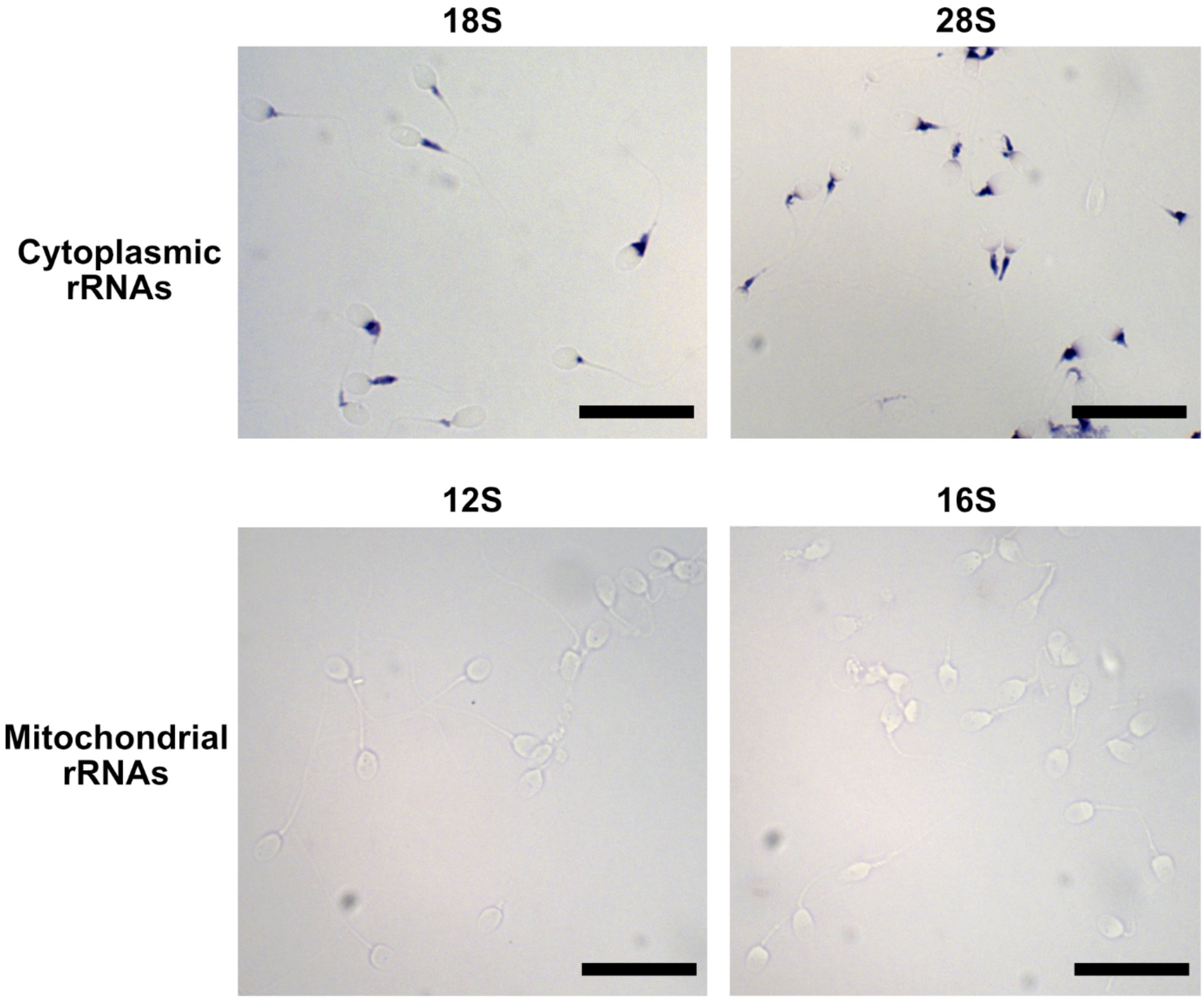


**Figure S3**. General views for the localization of rRNAs in human spermatozoa by *in situ* hybridization. Spermatozoa were labelled using antisense RNA probes targeting 28S, 18S, 16S, and 12S rRNAs. Scale bar: 10 μm.

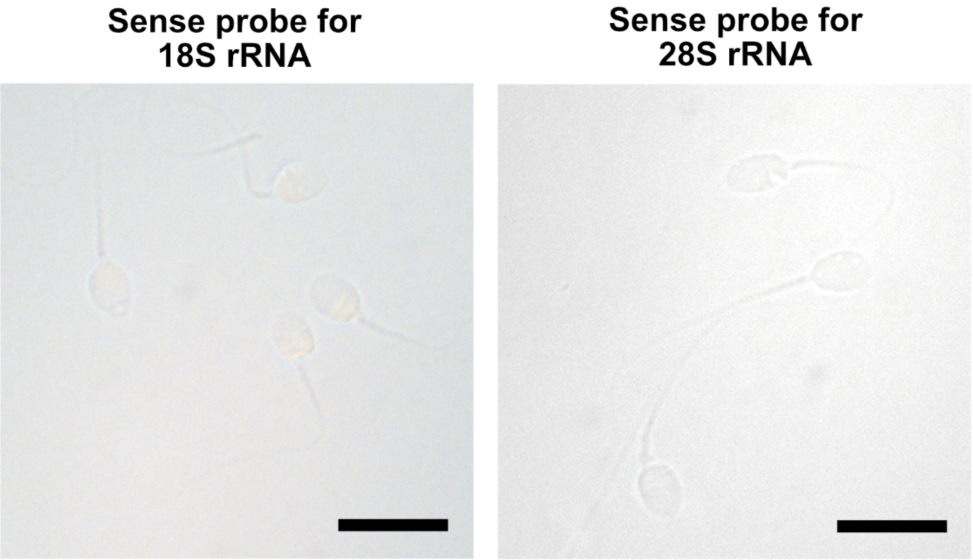


**Figure S4**. Negative controls for the localization of 18S and 28S rRNAs in human spermatozoa by *in situ* hybridization. Spermatozoa were incubated with sense RNA probes for 18S and 28S rRNAs. Scale bar: 10 µm.

**Table S1.** Ribosomal proteins in published human sperm proteomes → Excel file

**Table S2.** Primers used to synthesize probes for *in situ* hybridization

| Targeted RNA | Primers | Probe length (bp) |
|---|---|---|
| 28S | **CATTTAGGTGACACTATAGAAG**GCAGGAGGTGTCAGAAAAGTTACC<br>**GGATCCTAATACGACTCACTATAGG**TATTAGTGGGTGAACAATCCAACG | 150 |
| 18S | **CATTTAGGTGACACTATAGAAG**ACGATCAGATACCGTCGTAGTTCC<br>**GGATCCTAATACGACTCACTATAGG**CCTTTAAGTTTCAGCTTTGCAACC | 147 |
| 16S | **GGATCCTAATACGACTCACTATAGG**TGCAGAAGGTATAGGGGTTAGTCC | 151 |
| 12S | **GGATCCTAATACGACTCACTATAGG**GGGGTTTATCGATTACAGAACAGG | 157 |

In blue: promoter for the T7 RNA polymerase
In red: promoter for the sp6 RNA polymerase

**Table S3.** Number of individual sperm assessed for each condition (all replicates combined)

| **Parameter** | **CP (mg/ml)** | | | | **CHX (mg/ml)** | | | | | |
|---|---|---|---|---|---|---|---|---|---|---|
| | Control | 0.1 | 0.5 | 1.0 | Control | 0.1 | 0.5 | 1.0 | 1.5 | 2.0 |
| **Motile/Progressively motile sperm** | 1163 | 1007 | 924 | 1004 | 912 | 909 | 805 | 785 | 809 | 746 |
| **Live sperm (%)** | 2203 | 2129 | 2253 | 2178 | 2156 | | | 2168 | 2141 | 2181 |
| **VCL, VSL, VAP, ALHavg** | 870 | 754 | 553 | 428 | 701 | 531 | 443 | 435 | | 416 |
| **fBF, fAWL, P30** | 441 | 387 | 270 | 217 | 336 | 253 | 209 | 225 | | 201 |

ALHavg: average Amplitude of Lateral Head displacement; fAWL: flagellar arcwavelength; fBF: flagellar Beat Frequency; P30: Power of the first 30 µm of flagellum (1 fW = 1E-15 W); VAP: Average Path Velocity, VCL: Curvilinear Velocity, VSL: Straight Line Velocity.

**Table S4.** Transition list for the MRM analysis of selected proteins → Excel file

**Table S5.** Influence of mitochondrial (chloramphenicol, CP) and cytoplasmic (cycloheximide, CHX) ribosome inhibitors on sperm parameters, shown through fitting linear mixed-effects models. The fixed effect coefficients for intercept and concentration of inhibitors shown with 95% confidence intervals and number of donors n.

| Parameter | CP | | | | | CHX | | | | |
|---|---|---|---|---|---|---|---|---|---|---|
| | Intercept (95% CI) | p-value | Concentration (95% CI) | p-value | n | Intercept (95% CI) | p-value | Concentration (95% CI) | p-value | n |
| **Motile sperm (%)†** | 1.1 (0.80, 1.5) | <0.001** | -0.44 (-0.74, -0.14) | 0.0043* | 8 | 1.2 (0.91, 1.6) | <0.001** | -0.032 (-0.19, 0.12) | 0.68 | 7 |
| **Progressively motile sperm (%)†** | 0.88 (0.52, 1.2) | <0.001** | -0.80 (-1.2, -0.42) | <0.001** | 8 | 0.93 (0.61, 1.3) | <0.001** | -0.073 (-0.22 0.75) | 0.33 | 7 |
| **Live sperm (%)†** | 2.0 (1.7, 2.2) | <0.001** | -0.013 (-0.12, 0.099) | 0.82 | 7 | 2.0 (1.7, 2.2) | <0.001** | -0.029 (-0.11, 0.055) | 0.50 | 7 |
| **ATP (nM/1 million sperm)†** | 650 (570, 720) | <0.001** | -35 (-180, 110) | 0.63 | 9 | 660 (610, 710) | <0.001** | -31 (-80, 19) | 0.22 | 9 |
| **Relative densitometry phosphotyrosines/beta-tubulin†** | 1.1 (0.81, 1.4) | <0.001** | -0.011 (-0.44, 0.22) | 0.50 | 9 | 1.3 (0.81, 1.8) | <0.001** | -0.15 (-0.34, 0.029) | 0.10 | 7 |
| **VCL (µm/s)** | 98 (84, 110) | <0.001** | -28 (-41, -14) | <0.001** | 7 | 99 (87, 110) | <0.001** | 0.15 (-3.3, 3.6) | 0.93 | 7 |
| **VSL (µm/s)** | 58 (48, 68) | <0.001** | -15 (-22, -7.5) | <0.001** | 7 | 52 (42, 62) | <0.001** | -1.9 (-4.1, 0.27) | 0.086 | 7 |
| **VAP (µm/s)** | 49 (44, 54) | <0.001** | -8.8 (-13, -4.5) | <0.001** | 7 | †† | †† | †† | †† | 7 |
| **ALHavg (µm)** | 1.9 (1.5, 2.3) | <0.001** | -0.61 (-1.0, -0.21) | 0.0028* | 7 | 1.7 (1.4, 2.1) | <0.001** | 0.021 (-0.070, 0.11) | 0.66 | 7 |
| **fBF (µm)** | †† | †† | †† | †† | 7 | †† | †† | †† | †† | 7 |
| **fAWL (µm)** | 15 (14, 16) | <0.001** | -0.20 (-1.3, 0.86) | 0.71 | 7 | 16 (15, 17) | <0.001** | -0.075 (-0.55, 0.40) | 0.75 | 7 |
| **P30 (fW)** | 6.6 (5.2, 8.1) | <0.001** | -2.5 (-4.2, -0.82) | 0.0037* | 7 | 7.0 (6.0, 8.1) | <0.001** | 0.0064 (-1.3, 1.5) | 0.93 | 7 |

Purified human spermatozoa were incubated for 4 h in a capacitation medium in the absence (control) or presence of CP and CHX. For CP, all parameters were tested at concentrations 0.1, 0.5, and 1.0 mg/ml. For CHX, sperm motility, ATP content, and phosphotyrosine content were tested at concentrations 0.1, 0.5, 1.0, 1.5, and 2.0 mg/ml, vitality was measured at concentrations 1.0, 1.5, and 2.0 mg/ml, and kinematic parameters were tested at 0.1, 0.5, 1.0, and 2.0 mg/ml. For each parameter (P) a linear mixed effects model: P ~ 1 + Concentration + (1 + Concentration | Donor) was fit and the fixed effect coefficient for concentration reported. Models indicated by †† did not converge and the results are therefore not shown. * p-value ≤ 0.05, ** p-value ≤ 0.001. ALHavg: average Amplitude of Lateral Head displacement; fAWL: flagellar arcwavelength; fBF: flagellar Beat Frequency; P30: Power of the first 30 µm of flagellum (1 fW = 1E-15 W); VAP: Average Path Velocity, VCL: Curvilinear Velocity, VSL: Straight Line Velocity.

**Table S6**. Normalized abundance of proteins identified in CHX (Table S6a) and in CP (Table S6b) conditions → Excel file

**Table S7**. Fold change for each protein identified in CHX (Table S7a) and in CP (Table S7b) conditions → Excel file

**Table S8**. Raw data from the MRM analysis → Excel file
